# Effects of the collagen–glycosaminoglycan mesh on gene expression as determined by using principal component analysis-based unsupervised feature extraction

**DOI:** 10.1101/2021.09.20.461131

**Authors:** Y-h. Taguchi, Turki Turki

## Abstract

Development of the medical applications for substances or materials that contact the cells is important. Hence, it is necessary to elucidate how substance that surround cells affect the gene expression during incubation. Here, we compared the gene expression profiles of cell lines that were in contact with the collagen–glycosaminoglycan mesh and control cells. Principal component analysis-based unsupervised feature extraction was applied to identify genes with altered expression during incubation in the treated cell lines but not in the controls. The identified genes were enriched in various biological terms. Our method also outperformed a conventional methodology, namely, gene selection based on linear regression with time course.

## 1. Introduction

Several factors are known to affect cell division; one such effective factor is the contact with solid materials (or substance) [1]. Regulating the cell division process using biomaterials is the central theme of tissue engineering. The effect of tissue engineering scaffold is especially important because tissue engineering cannot be conducted without equipment that can store cell lines. Collagen–glycosaminoglycan mesh is one such important biomaterial because it is used to aid wound healing [2]. Although Klappericha and Bertozzi [3] once investigated the effect of collagen–glycosaminoglycan mesh on cell division cycles using microarray analysis, the small number of samples studied prevented them from identifying the genes whose expression significantly varied during development and whose expression profiles were distinct between controls and treated cells. Although they selected genes associated with *P*-values less than 0.001, considering the number of genes as 10^4^, it is far below significant.

The recently proposed principal component analysis (PCA)-based unsupervised feature extraction (FE) [4] has the ability to identify genes with expression profiles that were significantly different using small number of samples. In this study, we successfully applied PCA-based unsupervised FE to determine the gene expression profiles during cell division of cells in the control conditions and in contact with collagen–glycosaminoglycan mesh. The identified genes were found to be associated with several enrichment terms with considerable biological significance.

## 2. Materials and Methods

### 2.1. Gene expression profiles

Gene expression profiles were downloaded from the Gene Expression Omnibus (GEO) database (GEO ID: GSE6432). The dataset in GSE6432_series_matrix.txt.gz is available in the Series Matrix File(s) section. It consists of 32 gene expression profiles of the IMR90 cell lines, and the relevant details are provided in Table 1.

**Table 1.** The number of samples with the gene expression profiles. Treated means contact with the collagen–glycosaminoglycan mesh.

| Conditions | 1 hr | 2 hrs | 4 hrs | 8 hrs | 12 hrs | 24 hrs | 48 hrs | total |
| --- | --- | --- | --- | --- | --- | --- | --- | --- |
| treated | 3 | 2 | 3 | 4 | 3 | 2 | 2 | 10 |
| control | 1 | 2 | 3 | 2 | 3 | 1 | 1 | 13 |

### 2.2. PCA-based unsupervised FE

Gene expression profiles are formatted as matrices *x*_*ij*_ ∈ ℝ^22283×19^ for treated cells and *x*_*ij*_ ∈ ℝ^22283×13^ for control cells, where *x*_*ij*_ denotes gene expression of the *i*th probe at the *j*th sample. Before applying singular value decomposition (SVD), they were standardized as

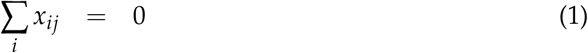

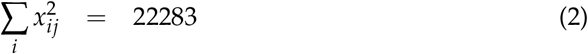

After applying SVD, we got the left hand singular value vector *u*_*ℓi*_, which corresponded to the principal component score attributed to the probes, and right hand singular value vector *v*_*ℓj*_, which corresponded to the principal component loadings attributed to the samples, if we interpreted the application of SVD as PCA.

In order to see which *v*_*ℓj*_ is coincident with time points, we applied linear regression as:

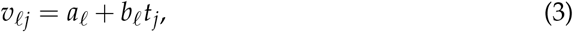

where *a*_*ℓ*_ and *b*_*ℓ*_ are regression coefficients and *t*_*j*_ is the time point (hrs in Table 1) associated with the *j*th sample. We used the `lm` function in R [5], and the obtained *P*-values were corrected using the Benjamini–Hochberg criterion [4]. *v*_3*j*_ for treated cell is associated with the adjusted *P*-values less than 0.05, whereas *v*_*ℓj*_s for control cell is not associated with adjusted *P*-values less than 0.05. This result is appropriate because the simple cell division process may not be associated with any time development other than cell senescence [6], which might not be detected in only 24 hours.

Probes are selected by assuming that *u*_3*i*_, associated with *v*_3*j*_, obeys the Gaussian distribution (null hypothesis) by assigning *P*-values to probes as:

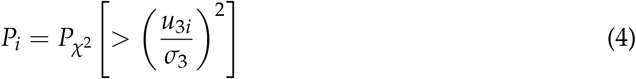

where 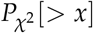 is the cumulative *χ*^2^ distribution, the argument is larger than *x*, and *σ*_3_ is the standard deviation. Thus, 324 probes associated with the adjusted *P*-value less than 0.01 were selected for the treated cell lines.

### 2.3. Gene selection using linear regression

As an alternative method to PCA-based unsupervised FE, we utilized linear regression-based FE. Linear regression is applied to *x*_*ij*_ as

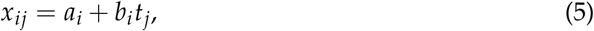

where *a*_*i*_ and *b*_*i*_ are regression coefficients and *t*_*j*_ is the time point (hrs in Table 1) associated with the *j*th sample. Subsequently, *i*s associated with adjusted *P*-values less than 0.01 were selected. The number of probes selected for treated cell lines was 813, and no probes were selected for the control cell lines.

### 2.4. Enrichment analysis

The IDs of the selected probes were converted to gene symbol using the ID converter in DAVID [7]. Then, the gene symbols converted from the prob IDs were uploaded to Enrichr [8].

## 3. Results

As mentioned in the Materials and Methods section, genes associated with the 318 probes for the treated cell lines (contact with collagen–glycosaminoglycan mesh) were uploaded to Enrichr (no probes were selected for control cell lines using this method). The full list of probes, genes and enrichment analysis is provided in the supplementary material (Data S1). Several enriched biological terms were determined.

The top ranked term in GO biological process (BP) (Table 2) is “regulation of apoptotic process”. Na et al reported [9] that collagen–glycosaminoglycan has an anti-apoptosis effect. Thus, the fact that this term is ranked first is reasonable.

**Table 2.**
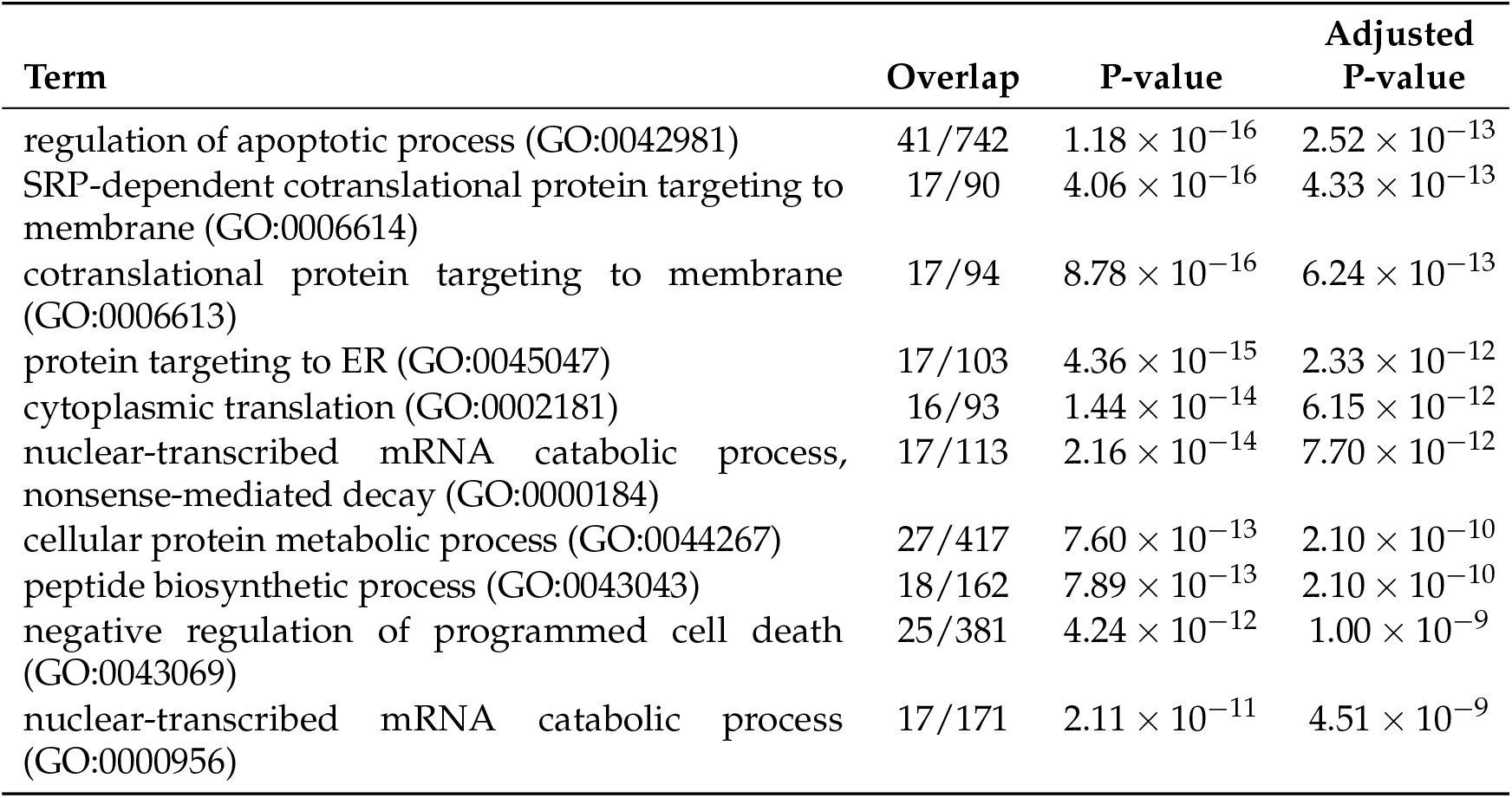
The top 10 enriched terms in “GO Biological Process 2021” using Enrichr. Overlap is the number of genes common between the genes uploaded and the genes in the category divided by the number of genes in the category. Probes, whose associated genes were uploaded to Enrichr, were identified using PCA-based unsupervised FE.

| Term | Overlap | P-value | Adjusted P-value |
| --- | --- | --- | --- |
| regulation of apoptotic process (GO:0042981) | 41/742 | $1.18 \times 10^{-16}$ | $2.52 \times 10^{-13}$ |
| SRP-dependent cotranslational protein targeting to membrane (GO:0006614) | 17/90 | $4.06 \times 10^{-16}$ | $4.33 \times 10^{-13}$ |
| cotranslational protein targeting to membrane (GO:0006613) | 17/94 | $8.78 \times 10^{-16}$ | $6.24 \times 10^{-13}$ |
| protein targeting to ER (GO:0045047) | 17/103 | $4.36 \times 10^{-15}$ | $2.33 \times 10^{-12}$ |
| cytoplasmic translation (GO:0002181) | 16/93 | $1.44 \times 10^{-14}$ | $6.15 \times 10^{-12}$ |
| nuclear-transcribed mRNA catabolic process, nonsense-mediated decay (GO:0000184) | 17/113 | $2.16 \times 10^{-14}$ | $7.70 \times 10^{-12}$ |
| cellular protein metabolic process (GO:0044267) | 27/417 | $7.60 \times 10^{-13}$ | $2.10 \times 10^{-10}$ |
| peptide biosynthetic process (GO:0043043) | 18/162 | $7.89 \times 10^{-13}$ | $2.10 \times 10^{-10}$ |
| negative regulation of programmed cell death (GO:0043069) | 25/381 | $4.24 \times 10^{-12}$ | $1.00 \times 10^{-9}$ |
| nuclear-transcribed mRNA catabolic process (GO:0000956) | 17/171 | $2.11 \times 10^{-11}$ | $4.51 \times 10^{-9}$ |

“Focal adhesion” is the top ranked term in “GO Cellular Component 2021” (Table 3); moreover, Murphy et al [10] reported that the collagen–glycosaminoglycan scaffold plays critical roles in focal adhesion.

**Table 3.** The top 10 enriched terms in “GO Cellular Component 2021” using Enrichr. Overlap is the number of genes common between the genes uploaded and the genes in the category divided by the number of genes in the category. Probes, whose associated genes were uploaded to Enrichr, were identified using PCA-based unsupervised FE.

| Term | Overlap | P-value | Adjusted P-value |
| --- | --- | --- | --- |
| focal adhesion (GO:0005925) | 43/387 | $2.22 \times 10^{-29}$ | $4.77 \times 10^{-27}$ |
| cell-substrate junction (GO:0030055) | 43/394 | $4.69 \times 10^{-29}$ | $5.04 \times 10^{-27}$ |
| intracellular organelle lumen (GO:0070013) | 40/848 | $5.63 \times 10^{-14}$ | $4.03 \times 10^{-12}$ |
| collagen-containing extracellular matrix (GO:0062023) | 25/380 | $4.00 \times 10^{-12}$ | $2.15 \times 10^{-10}$ |
| endoplasmic reticulum lumen (GO:0005788) | 21/285 | $2.78 \times 10^{-11}$ | $1.19 \times 10^{-9}$ |
| cytosolic large ribosomal subunit (GO:0022625) | 10/55 | $7.95 \times 10^{-10}$ | $2.85 \times 10^{-8}$ |
| large ribosomal subunit (GO:0015934) | 10/59 | $1.64 \times 10^{-9}$ | $5.03 \times 10^{-8}$ |
| ribosome (GO:0005840) | 10/62 | $2.72 \times 10^{-9}$ | $7.30 \times 10^{-8}$ |
| secretory granule lumen (GO:0034774) | 19/316 | $7.48 \times 10^{-9}$ | $1.79 \times 10^{-7}$ |
| ficolin-1-rich granule lumen (GO:1904813) | 12/123 | $2.50 \times 10^{-8}$ | $5.37 \times 10^{-7}$ |

Other than these three categories, there are some additional categories that support the suitability of our analysis. For example, “ARCHS4 Cell-lines” lists the IMR90, which is the cell line used in this study, as the top ranked cell line (Table 5).

**Table 4.** The top 10 enriched terms in “KEGG 2021 Human” using Enrichr. Overlap is the number of genes common between the genes uploaded and the genes in the category divided by the number of genes in the category. Probes, whose associated genes were uploaded to Enrichr, were identified using PCA-based unsupervised FE.

| Term | Overlap | P-value | Adjusted P-value |
| --- | --- | --- | --- |
| Coronavirus disease | 21/232 | $5.36 \times 10^{-13}$ | $1.22 \times 10^{-10}$ |
| Ribosome | 17/158 | $5.88 \times 10^{-12}$ | $6.67 \times 10^{-10}$ |
| Legionellosis | 9/57 | $2.09 \times 10^{-8}$ | $1.58 \times 10^{-6}$ |
| Salmonella infection | 16/249 | $4.75 \times 10^{-8}$ | $2.70 \times 10^{-6}$ |
| IL-17 signaling pathway | 10/94 | $1.64 \times 10^{-7}$ | $7.46 \times 10^{-6}$ |
| Glycolysis / Gluconeogenesis | 8/67 | $1.20 \times 10^{-6}$ | $4.52 \times 10^{-5}$ |
| Lipid and atherosclerosis | 13/215 | $1.75 \times 10^{-6}$ | $5.69 \times 10^{-5}$ |
| Protein digestion and absorption | 9/103 | $3.65 \times 10^{-6}$ | $1.04 \times 10^{-4}$ |
| Focal adhesion | 12/201 | $5.03 \times 10^{-6}$ | $1.17 \times 10^{-4}$ |
| PI3K-Akt signaling pathway | 16/354 | $5.14 \times 10^{-6}$ | $1.17 \times 10^{-4}$ |

**Table 5.** The top 10 enriched terms in “ARCHS4 Cell-lines” using Enrichr. Overlap is the number of genes common between the genes uploaded and the genes in the category divided by the number of genes in the category. Probes, whose associated genes were uploaded to Enrichr, were identified using PCA-based unsupervised FE.

| Term | Overlap | P-value | Adjusted P-value |
| --- | --- | --- | --- |
| IMR90 | 89/2395 | $1.56 \times 10^{-24}$ | $1.95 \times 10^{-22}$ |
| NHDF | 79/2395 | $2.70 \times 10^{-18}$ | $1.69 \times 10^{-16}$ |
| BJ CELL | 72/2395 | $2.02 \times 10^{-14}$ | $8.41 \times 10^{-13}$ |
| HUVEC | 64/2395 | $1.61 \times 10^{-10}$ | $5.04 \times 10^{-9}$ |
| T24 | 62/2395 | $1.23 \times 10^{-9}$ | $3.09 \times 10^{-8}$ |
| T98G | 59/2395 | $2.22 \times 10^{-8}$ | $4.63 \times 10^{-7}$ |
| BT549 | 56/2395 | $3.28 \times 10^{-7}$ | $5.12 \times 10^{-6}$ |
| DU145 | 56/2395 | $3.28 \times 10^{-7}$ | $5.12 \times 10^{-6}$ |
| CAKI1 | 55/2395 | $7.68 \times 10^{-7}$ | $9.60 \times 10^{-6}$ |
| U87 | 55/2395 | $7.68 \times 10^{-7}$ | $9.60 \times 10^{-6}$ |

Moreover, although it is not the top ranked term, “FETAL LUNG”, from which IMR90 cell lines were derived, is ranked within the top 10 ranked terms in “ARCHS4 Tissues” (Table 6).

**Table 6.**
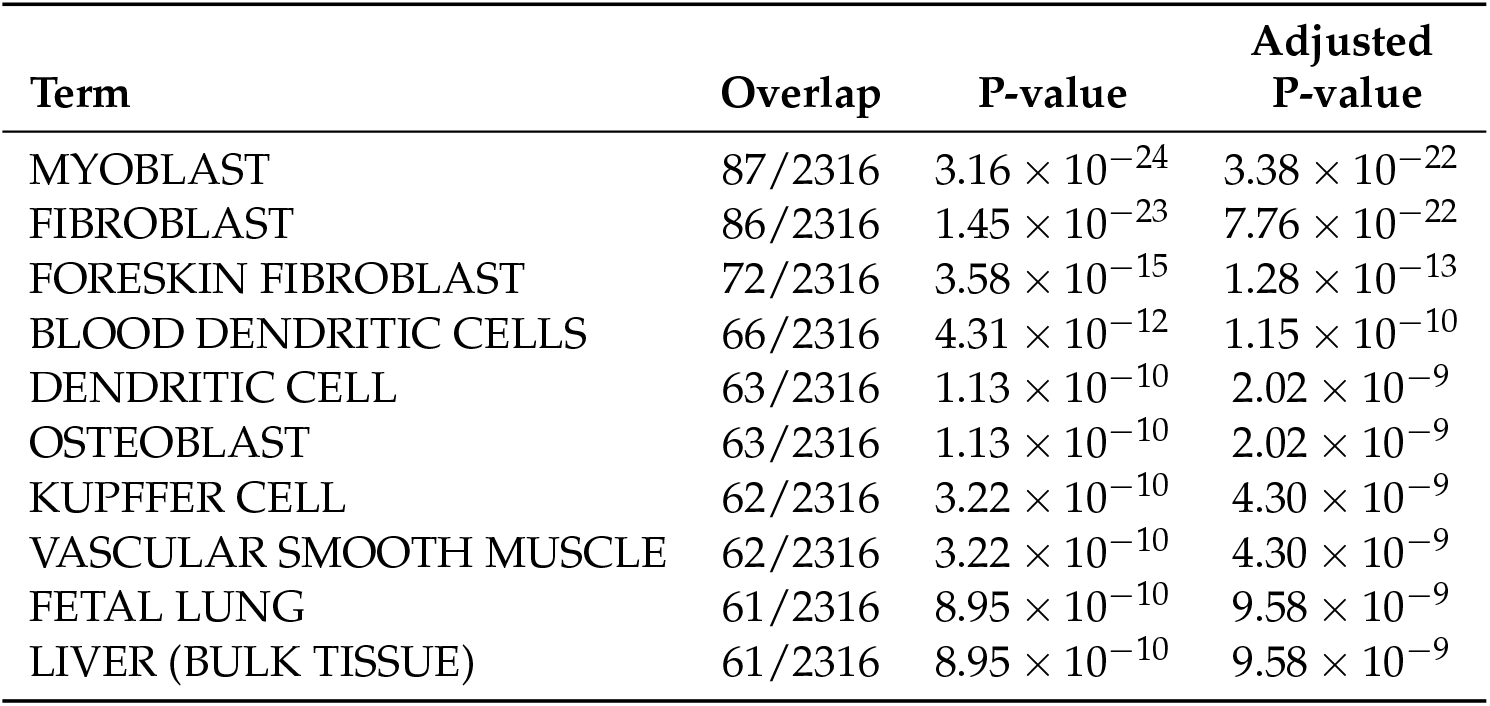
The top 10 enriched terms in “ARCHS4 Tissues” using Enrichr. Overlap is the number of genes common between the genes uploaded and the genes in the category divided by the number of genes in the category. Probes, whose associated genes were uploaded to Enrichr, were identified using PCA-based unsupervised FE.

Although we have provided only a few examples, our results suggest that our analysis was robust.

## 4. Discussion

Although we have successfully applied our methodology to the data set, one might wonder whether more conventional method can achieve similar performance. Since this dataset was generated using archaic technology, namely, microarray, more modernized methodologies adapted to high-throughput sequencing technology (e.g., edgeR [11] or DESeq2 [12]) cannot be employed. As for the archaic technologies adapted to microarray (e.g., SAM [13] and limma [14]) cannot be employed as well because they can deal with only categorical classification, whereas we need to identify genes whose expression is altered as a numerical variable (hours). Thus, we decided to employ more a more conventional methodology than SAM or limma, namely, gene selection using linear regression.

As described in the Materials and Methods section, we identified 813 probes using linear regression-based FE and uploaded the gene symbols associated with the identified probes to Enrichr. When considering only the number of probes selected, it was better than the PCA-based unsupervised FE, which could identify only 324 probes. Selecting no probes for the control cell lines is the same as PCA-based unsupervised FE. Thus, it seems that the application of PCA-based unsupervised FE, instead of linear regression, was not productive.

Nevertheless, if we consider the performance of the enrichment analysis more carefully, this impression is reversed. Full list of probes, genes, and the results of enrichment analysis are provided in the supplementary material (Data S2). First, for “GO BP 2021”, in which PCA-based unsupervised FE ranked apoptosis first (Tables 2, and 7), although the top ranked term “regulation of apoptotic process” in Table 2 is associated with adjusted *P*-value as small as 2.52 × 10^−13^, the top ranked term in Table 7 is associated with adjusted *P*-value as large as 4.56 × 10^−2^, which is much less significant. Even the tenth ranked term in Table 2 is more significant than the top ranked term in Table 7. Generally, more genes uploaded have more opportunity to be associated with more significant enrichment. Nevertheless, genes associated with 813 probes, which is greater than the 324 probes identified using PCA-based unsupervised FE, could be associated with the less significant terms. This definitely suggests the inferiority of linear regression as compared to PCA-based unsupervised FE.

**Table 7.** The top 10 enriched terms in “GO Biological Process 2021” using Enrichr. Overlap is the number of genes common between the genes uploaded and the genes in the category divided by the number of genes in the category. Probes, whose associated genes were uploaded to Enrichr, were identified using linear regression.

| Term | Overlap | P-value | Adjusted P-value |
| --- | --- | --- | --- |
| actin polymerization or depolymerization (GO:0008154) | 9/50 | $4.05 \times 10^{-5}$ | $4.56 \times 10^{-2}$ |
| rRNA-containing ribonucleoprotein complex export from nucleus (GO:0071428) | 4/7 | $4.22 \times 10^{-5}$ | $4.56 \times 10^{-2}$ |
| positive regulation of protein modification process (GO:0031401) | 20/214 | $4.32 \times 10^{-5}$ | $4.56 \times 10^{-2}$ |
| transmembrane receptor protein tyrosine kinase signaling pathway (GO:0007169) | 30/404 | $5.44 \times 10^{-5}$ | $4.56 \times 10^{-2}$ |
| protein stabilization (GO:0050821) | 17/179 | $1.31 \times 10^{-4}$ | $7.08 \times 10^{-2}$ |
| regulation of lipid biosynthetic process (GO:0046890) | 7/35 | $1.46 \times 10^{-4}$ | $7.08 \times 10^{-2}$ |
| regulation of cellular metabolic process (GO:0031323) | 8/47 | $1.63 \times 10^{-4}$ | $7.08 \times 10^{-2}$ |
| positive regulation of cellular protein metabolic process (GO:0032270) | 12/102 | $1.74 \times 10^{-4}$ | $7.08 \times 10^{-2}$ |
| regulation of stress fiber assembly (GO:0051492) | 10/74 | $1.90 \times 10^{-4}$ | $7.08 \times 10^{-2}$ |
| regulation of mRNA catabolic process (GO:0061013) | 13/122 | $2.58 \times 10^{-4}$ | $7.65 \times 10^{-2}$ |

As for the comparison of the “GO Cellular Component 2021” between Tables 3 and 8, we have a similar impression.

**Table 8.** The top 10 enriched terms in “GO Cellular Component 2021” using Enrichr. Overlap is the number of genes common between the genes uploaded and the genes in the category divided by the number of genes in the category. Probes, whose associated genes were uploaded to Enrichr, were identified using linear regression.

| Term | Overlap | P-value | Adjusted P-value |
| --- | --- | --- | --- |
| focal adhesion (GO:0005925) | 39/387 | $1.41 \times 10^{-9}$ | $3.39 \times 10^{-7}$ |
| cell-substrate junction (GO:0030055) | 39/394 | $2.34 \times 10^{-9}$ | $3.39 \times 10^{-7}$ |
| actin cytoskeleton (GO:0015629) | 25/316 | $8.27 \times 10^{-5}$ | $7.97 \times 10^{-3}$ |
| intracellular organelle lumen (GO:0070013) | 49/848 | $2.02 \times 10^{-4}$ | $1.46 \times 10^{-2}$ |
| nucleus (GO:0005634) | 186/4484 | $1.06 \times 10^{-3}$ | $5.51 \times 10^{-2}$ |
| collagen-containing extracellular matrix (GO:0062023) | 25/380 | $1.30 \times 10^{-3}$ | $5.51 \times 10^{-2}$ |
| cytoplasmic stress granule (GO:0010494) | 8/65 | $1.54 \times 10^{-3}$ | $5.51 \times 10^{-2}$ |
| intracellular membrane-bounded organelle (GO:0043231) | 210/5192 | $1.66 \times 10^{-3}$ | $5.51 \times 10^{-2}$ |
| endoplasmic reticulum lumen (GO:0005788) | 20/285 | $1.81 \times 10^{-3}$ | $5.51 \times 10^{-2}$ |
| intracellular non-membrane-bounded organelle (GO:0043232) | 58/1158 | $1.91 \times 10^{-3}$ | $5.51 \times 10^{-2}$ |

Although “focal adhesion” is ranked first in both the Tables, its significance is very distinct. It is associated with adjusted *P*-value as small as 4.77 × 10^−27^ in Table 3 whereas it is associated with that as large as 3.39 × 10^−7^ in Table 8. The number of overlapping gene is only 39 in Table 8 whereas it is more (43) in Table 3, despite the fact that more total number of genes was uploaded to Enrichr as shown in Table 8. Thus, the performance of linear regression is again poorer than that of PCA-based unsupervised FE.

For KEGG, not only the generally adjusted *P*-values are larger (i.e. less significant) in Table 9 than those in Table 4, but also neither “Glycolysis / Gluconeogenesis” nor “Focal adhesion”, which are ranked within the top 10 in Table 4, are even not listed in Table 9 and no other terms seemingly related to the experiments are mentioned. Thus, the performance of linear regression is again poorer than that of PCA-based unsupervised FE.

**Table 9.** The top 10 enriched terms in “KEGG 2021 Human” using Enrichr. Overlap is the number of genes common between the genes uploaded and the genes in the category divided by the number of genes in the category. Probes, whose associated genes were uploaded to Enrichr, were identified using linear regression.

| Term | Overlap | P-value | Adjusted P-value |
| --- | --- | --- | --- |
| PI3K-Akt signaling pathway | 28/354 | $3.17 \times 10^{-5}$ | $9.30 \times 10^{-3}$ |
| Sphingolipid signaling pathway | 13/119 | $2.01 \times 10^{-4}$ | $1.66 \times 10^{-2}$ |
| Arrhythmogenic right ventricular cardiomyopathy | 10/77 | $2.65 \times 10^{-4}$ | $1.66 \times 10^{-2}$ |
| Antigen processing and presentation | 10/78 | $2.95 \times 10^{-4}$ | $1.66 \times 10^{-2}$ |
| Hepatitis C | 15/157 | $2.99 \times 10^{-4}$ | $1.66 \times 10^{-2}$ |
| Salmonella infection | 20/249 | $3.39 \times 10^{-4}$ | $1.66 \times 10^{-2}$ |
| Hippo signaling pathway | 15/163 | $4.47 \times 10^{-4}$ | $1.87 \times 10^{-2}$ |
| Tight junction | 15/169 | $6.54 \times 10^{-4}$ | $2.29 \times 10^{-2}$ |
| Protein processing in endoplasmic reticulum | 15/171 | $7.39 \times 10^{-4}$ | $2.29 \times 10^{-2}$ |
| AMPK signaling pathway | 12/120 | $7.81 \times 10^{-4}$ | $2.29 \times 10^{-2}$ |

For “ARCHS4 Cell-lines” and “ARCHS4 Tissue”, the results are similar. In Table 10, not only the adjusted *P*-values generally larger (i.e. less significant) than those in Table 5 but also the adjusted *P*-values attributed to IMR90 in Table 10 (1.06 × 10^−5^) are much larger (i.e. less significant) than those in Table 5. The number of overlapping genes for IMR90 is only 128 in Table 5 whereas that in Table 10 is 89, despite of the fact that more than twice the total number of genes were uploaded to Enrichr, as shown in Table 5. However, the number of overlapping genes for HUVEC, which is the wrong one, is as large as 113 in Table 10, whereas that in Table 5 is only 64. Thus, the increased number of genes selected using linear regression contributes substantially to the increase in overlapping genes assigned to the wrong answer. Moreover, lower ranked terms failed to demonstrate an association with significant *P*-values (e.g. less than 0.015). These finding suggest the inferiority of linear regression as compared to PCA-based unsupervised FE.

**Table 10.** The top 10 enriched terms in “ARCHS4 Cell-lines” using Enrichr. Overlap is the number of genes common between the genes uploaded and the genes in the category divided by the number of genes in the category. Probes, whose associated genes were uploaded to Enrichr, were identified using linear regression.

| Term | Overlap | P-value | Adjusted P-value |
| --- | --- | --- | --- |
| IMR90 | 128/2395 | $8.49 \times 10^{-8}$ | $1.06 \times 10^{-5}$ |
| HUVEC | 113/2395 | $1.54 \times 10^{-4}$ | $9.65 \times 10^{-3}$ |
| NHDF | 112/2395 | $2.35 \times 10^{-4}$ | $9.78 \times 10^{-3}$ |
| BT549 | 103/2395 | $6.32 \times 10^{-3}$ | $1.62 \times 10^{-1}$ |
| BJ CELL | 101/2395 | $1.17 \times 10^{-2}$ | $1.62 \times 10^{-1}$ |
| HNSCC | 101/2395 | $1.17 \times 10^{-2}$ | $1.62 \times 10^{-1}$ |
| KNS42 | 101/2395 | $1.17 \times 10^{-2}$ | $1.62 \times 10^{-1}$ |
| NHBE | 101/2395 | $1.17 \times 10^{-2}$ | $1.62 \times 10^{-1}$ |
| U87 | 101/2395 | $1.17 \times 10^{-2}$ | $1.62 \times 10^{-1}$ |
| DAOY | 99/2395 | $2.07 \times 10^{-2}$ | $2.35 \times 10^{-1}$ |

Although “FETAL LUNG” is fourth ranked in Table 11, adjusted *P*-value is 1.05 × 10^−3^, which is much less significant than that in Table 6 (9.58 × 10^−9^). Thus, overall, PCA-based unsupervised FE performed better than linear regression.

**Table 11.** The top 10 enriched terms in “ARCHS4 Tissues” using Enrichr. Overlap is the number of genes common between the genes uploaded and the genes in the category divided by the number of genes in the category. Probes, whose associated genes were uploaded to Enrichr, were identified using linear regression.

| Term | Overlap | P-value | Adjusted P-value |
| --- | --- | --- | --- |
| FIBROBLAST | 116/2316 | $9.27 \times 10^{-6}$ | $5.01 \times 10^{-4}$ |
| VENTRICLE | 116/2316 | $9.27 \times 10^{-6}$ | $5.01 \times 10^{-4}$ |
| ADIPOSE (BULK TISSUE) | 113/2316 | $3.89 \times 10^{-5}$ | $1.05 \times 10^{-3}$ |
| FETAL LUNG | 113/2316 | $3.89 \times 10^{-5}$ | $1.05 \times 10^{-3}$ |
| RESPIRATORY SMOOTH MUSCLE | 112/2316 | $6.14 \times 10^{-5}$ | $1.33 \times 10^{-3}$ |
| OMENTUM | 111/2316 | $9.58 \times 10^{-5}$ | $1.73 \times 10^{-3}$ |
| SUBCUTANEOUS ADIPOSE TISSUE | 106/2316 | $7.60 \times 10^{-4}$ | $1.17 \times 10^{-2}$ |
| MYOBLAST | 104/2316 | $1.61 \times 10^{-3}$ | $2.18 \times 10^{-2}$ |
| NEURONAL EPITHELIUM | 103/2316 | $2.31 \times 10^{-3}$ | $2.77 \times 10^{-2}$ |
| ASTROCYTE | 99/2316 | $8.75 \times 10^{-3}$ | $8.54 \times 10^{-2}$ |

## 5. Conclusions

Here, we applied PCA-based unsupervised FE to gene expression profiles for the IMR90 cell lines incubated in collagen–glycosaminoglycan mesh. Whereas no genes whose expression vary as time goes are detected in control cell lines, the expression profiles of several genes were altered during the cell division process. These genes are associated with several enriched biological terms. One conventional method, linear regression was employed for comparison. Although it could select several hundred genes whose expression vary as time passes, their enrichment was inferior to that seen using PCA-based unsupervised FE. Thus, PCA-based unsupervised FE can not only achieve good performance but also outperform a conventional method.

## Author Contributions

YHT planned the research, performed analyses. YHT and TT have evaluated the results, discussions, outcomes and wrote and reviewed the manuscript.

## Data Availability Statement

Data used in this study is available in GEO ID GSE6432.

